# The olfactory organ is a unique site for resident neutrophils in the brain

**DOI:** 10.1101/2021.07.22.453396

**Authors:** M. Fernanda Palominos, Danissa Candia, Jorge Torres Paz, Kathleen E. Whitlock

## Abstract

For decades we have known that the brain “drains” through the subarachnoid space following a route that crosses the cribriform plate to the nasal mucosa and cervical lymph nodes. Yet little is known about the potential role of the olfactory epithelia and associated lymphatic vasculature in the immune response. To better understand the immune response in the olfactory organs we used cell-specific fluorescent reporter lines in dissected, intact adult brains to visualize blood-lymphatic vasculature and neutrophils in the olfactory sensory system. Here we show that the extensive blood vasculature of the olfactory organs is associated with a lymphatic cell type resembling high endothelial venules (HEVs) of the lymph nodes in mammals and a second resembling Mural Lymphatic Endothelial Cells (muLECs) that extended from the brain to the peripheral olfactory epithelia. Surprisingly, the olfactory organs contained the only neutrophil populations observed in the brain. Damage to the olfactory epithelia resulted in a rapid increase of neutrophils within the olfactory organs as well as the appearance of neutrophils in the brain suggesting that neutrophils enter the brain in response to damage. Analysis of cell division during and after damage showed an increase in BrdU labeling in the olfactory epithelia and a subset of the neutrophils. Our results reveal a unique population of neutrophils in the olfactory organs that are associated with an extensive lymphatic vasculature suggesting a dual olfactory-immune function for this unique sensory system.

**Highlights:**

- The olfactory organ is the only region of the brain that contains resident neutrophils in the adult animal.
- Damage to olfactory sensory neurons triggers a rapid mobilization of neutrophils within the olfactory organ and in the central nervous system.
- Two types of lymphatic vasculature resembling Mural Lymphatic Endothelial Cells (muLEC) and High Endothelial Venules (HEV) are present in the olfactory sensory system.
- Lymphatic vasculature resembling Mural Lymphatic Endothelial Cells (muLEC) wrap the olfactory bulbs and extend across the cribriform plate to olfactory epithelia.

## Introduction

### The adult olfactory organ blood-lymphatic system

In vertebrates the olfactory sensory neurons (OSNs), a group of continually renewing neurons located in the olfactory epithelium (OE), extend their axons across the cribriform plate where they make their first synapses in the olfactory bulb (OB) (Sakano, 2010; Whitlock, 2015). This connection between the OE and the OB is part of a complex neural and immune interface that includes flow of cerebral spinal fluid (CSF) and interstitial fluid (ISF) from the subarachnoid space toward the nasal mucosa. Evidence supporting a connection between the subarachnoid space of the brain and cervical lymph nodes via the nasal mucosa was first proposed over a century ago (for review see: (Faber, 1937; Jackson *et al*., 1979). Subsequent studies in mammals using labeled tracers confirmed a drainage route from the cranial subarachnoid space through the olfactory pathway leaving the nasal mucosa via terminal lymphatics or into blood capillaries (Cserr *et al*., 1992). Thus the potential for turnover of brain extracellular fluids, via drainage to blood and deep cervical lymph, presented a system whereby immunogenic material and immune cells from the central nervous system (CNS) could pass to immune organs outside the brain via the olfactory epithelia.

The lymphatic system of vertebrates, composed of lymphatic vessels, lymphoid organs/tissues and the circulating lymph fluid, is highly conserved at the functional level (Boehm *et al*., 2012) and is suggested to have originated in teleost fishes where the heart provided the energy to propel lymph through vessels associated with the primary vasculature (Hedrick *et al*., 2013). Lymphocytes are generated in primary lymphoid organs (thymus and bone marrow: mammals / thymus and kidney; teleost fishes) and maintained in secondary lymphoid tissues (spleen and lymph nodes; mammals/spleen, nasopharynx and gill tissues: teleost fishes) (Bjørgen and Koppang, 2021). Of particular interest are the nasopharynx-associated lymphoid tissues (NALT), a term used in mammals to describe the network of lymphoid tissue in the pharynx and palate (tonsils). Teleost fish lack organized lymphoid structures such as tonsils yet a recent study suggested the presence of a NALT-like diffuse network of lymphoid and myeloid cells scattered both intraepithelial and in the lamina propria of the fish olfactory organ (Tacchi *et al*., 2014).

More recently the “re-discovery” of lymphatic vasculature associated with the meninges in the central nervous system (CNS) of mammals (Aspelund *et al*., 2015);(Louveau *et al*., 2015);(Da Mesquita *et al*., 2018);(Dolgin, 2020) and of zebrafish (Bower *et al*., 2017); (Bower and Hogan, 2018) coupled with studies indicating that the meninges contain a diverse array of immune cells that can migrate via the sinus-associated meningeal lymphatic vessels and/or via cribriform plate and nasal lymphatics into cervical lymph nodes (Rua and McGavern, 2018)(Rustenhoven *et al*., 2021) (Sun *et al*., 2018), have led to a renewed interest in immune trafficking in the nervous system. To date in spite of over a century of reports on “brain drainage” through the olfactory system/nasal mucosa and the expanded knowledge of lymphatic vasculature in the vertebrate brain, there are no detailed descriptions of the lymphatic vasculature (LV) in the olfactory organ.

### Neutrophils and the Nervous System

Neutrophils the most abundant type of white blood cells are normally found in the blood stream where they are rapidly recruited to a site of injury or infection and perform a critical role in inflammation and pathogen clearance. Neutrophils have been shown to interact with and regulate not only the innate but also the adaptive immune cells where they can rapidly migrate via afferent lymphatics of inflamed tissues to lymph nodes (Voisin and Nourshargh, 2019) (Beauvillain *et al*., 2011) (Hampton *et al*., 2015; Maletto *et al*., 2006). Thus neutrophils migrate not only on the blood vasculature and interstitial tissues, but can migrate into the lymphoid system and are in the unique position to participate in the very early stages of both innate and adaptive immune responses. Under normal conditions, neutrophils are scarce in the CNS where the brain–blood barrier (BBB) prevents their migration into the brain parenchyma and cerebrospinal fluid. Conditions of neuroinflammation and injury induced damage to the BBB are associated with the infiltration of the CNS by neutrophils (Harrison-Brown *et al*., 2016; Khorooshi *et al*., 2020; Manda-Handzlik and Demkow, 2019).

Previously, we performed both microarray and RNAseq analyses (Harden *et al*., 2006) (Calfun *et al*., 2016) of zebrafish adult OE to investigate differentially expressed genes involved in the formation of olfactory memory (Whitlock, 2006). In addition to known genes expressed in the OE, we found genes specific to both the innate and the adaptive immune systems (Calfún, 2017), prompting us to investigate the potential “immune architecture” of the OE.

We have shown that neutrophils populate the developing olfactory organ and use the blood vasculature to migrate to the olfactory organ in response to injury (Palominos and Whitlock, 2020). With the recent re-discovery of the CNS lymphatics in mammals and zebrafish (Bower and Hogan, 2018; Louveau *et al*., 2015), we examined the extent of lymphatic vasculature in the adult olfactory organs and its association with blood vasculature. Neutrophils, known to play a key role in both the innate and the adaptive immune response (Odobasic *et al*., 2016);(Meinderts *et al*., 2019);(Yang *et al*., 2010), were always found in the olfactory organ of adult zebrafish under both normal and damage conditions. In fishes, the olfactory bulb may be involved in immune responses where activation of immune cells in the olfactory bulb resulted from peripheral neuronal signals (Das and Salinas, 2020). Our results suggest that the olfactory organ has the potential to respond quickly to damage via a local population of neutrophils located in both the neuronal and non-neuronal tissues of the olfactory organ.

## RESULTS

### The Adult Olfactory Sensory System has Extensive Lymphatic Vasculature

Previously we have shown that the lymphatic vasculature (LV) associated with the developing olfactory organs is evident at 14 days post fertilization (dpf) initiating in the ventrolateral side of the organ (Palominos and Whitlock, 2020). To better understand the LV system in the olfactory sensory system of the adult we dissected brains, with olfactory organs attached, from *Tg(lyve1b:EGFP;OMP:RFP)* animals (Fig. 1). The olfactory organs (OO) are made up of sensory epithelia containing the OMP:RFP^+^ sensory neurons (Fig. 1A-D, F, red) and respiratory epithelia, surrounded by what appears to be an extension of the epineurium (EN) of the olfactory nerve (Fig 1A-D, EN). At this point it is not clear where the meningeal membranes fuse with the epineurium after crossing the cribriform plate (Jackson *et al*., 1979). Viewed from the dorsal side, lyve1b:EGFP^+^ LV were found in the OO (Fig 1A, OO, green, B, green, arrowheads), olfactory bulb (Fig 1A, OB, green, arrow) and diencephalon (Fig. 1A, TeO, green, arrow), but not the telencephalon. In the dorsal OO the lyve1b:EGFP^+^ cells (Fig. 1B, green, arrowheads) line the lamellae of the OE (Fig. 1B, LOE). In contrast, when viewed from the ventral side there was an apparently continuous network of LV extending from the OO to the OB and along ventral telencephalon (Fig. 1 C, D, green). The lyve1b:EGFP^+^ cells were also evident in the ventral OO associated with the olfactory nerve (Fig. 1D, ON, red). Two morphologically distinct lymphatic cell types were observed. In the OO thick tubular cells associated with the LOE (Fig. 1B, D, arrowheads), resembling High Endothelial Venules (HEV-like, HEV-L; Fig. 1E) that control lymphocyte trafficking in mammals (Ager, 2017). To date these cells have not been described in the peripheral olfactory sensory system. In the OB smaller lyve1b:EGFP^+^ cells covering the dorsal OB and ventral telencephalon, apparently connected by fine processes, resembled Mural Lymphatic Endothelial Cells (muLEC-L) after Bower (Bower *et al*., 2017) (Fig. 1D, arrows, green, F, muLEC-L, green). This cell type was also observed in the OO (Figs. 2, 3). In contrast to the cells described by Bower, the muLEC-L appeared to be connected by fine processes (Fig. 1, F, arrows) and not separate cells like the BV-associated muLECs (Bower *et al*., 2017). At this time it is not clear whether these connections have a lumen. Thus, in adult zebrafish there is an extensive LV system associated with the olfactory sensory system (Fig. 1) wrapping the OE (HEV-L), encompassing the olfactory bulb (muLEC-L) with apparently continuous connections along the ventral telencephalon (Fig. 1 C).

**Figure 1.**
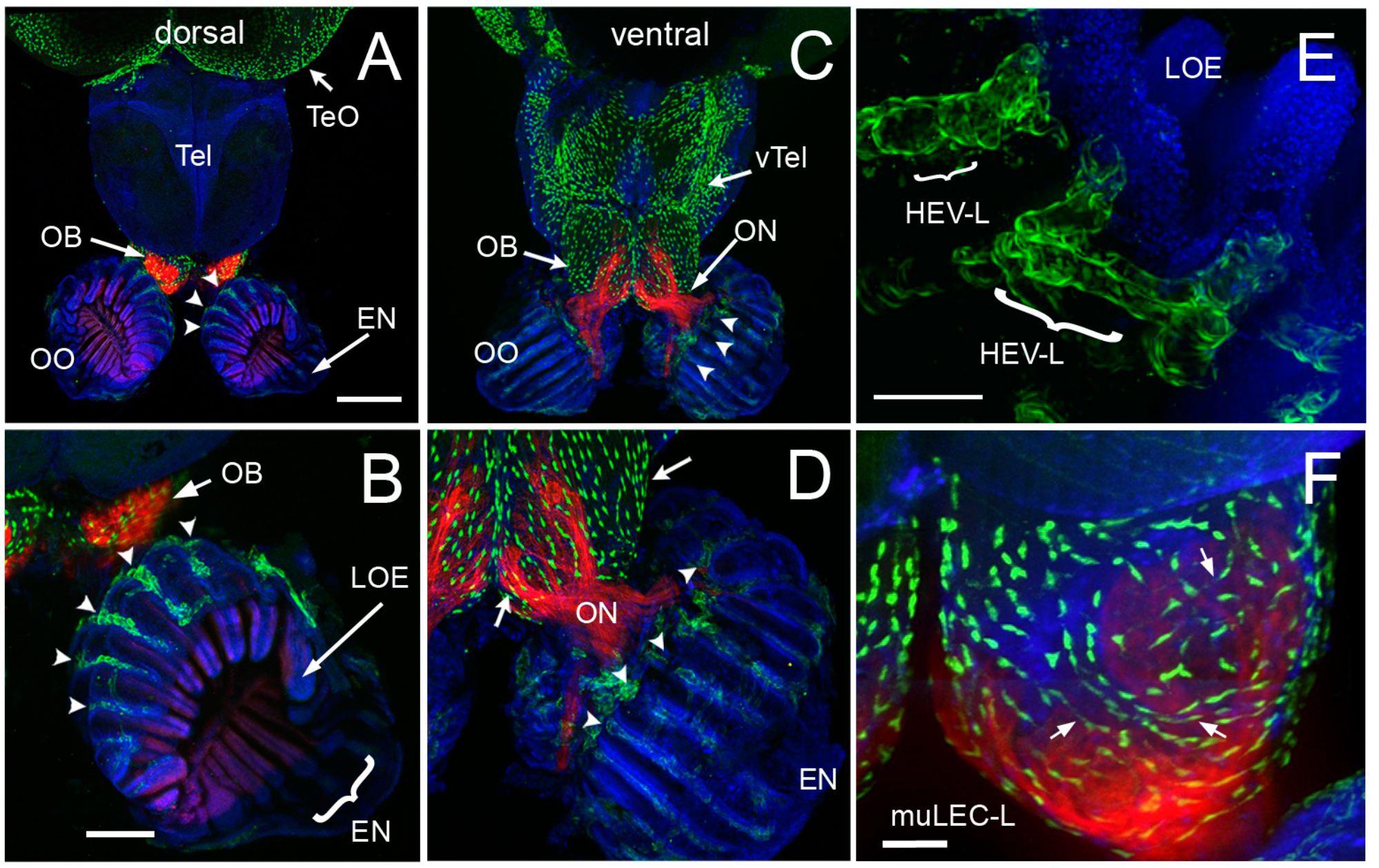
The adult olfactory organ have an extensive blood-lymphatic system. **A-F**. Whole mount brains of adult *lyve1b:EGFP;OMP:RFP* animals with OSN (red) and lymphatic vasculature (green). **A** The OE and OBs have extensive lymphatic vasculature (LV, green) but not dorsal the telencephalon (Tel). **B.** Higher magnification of OO in A with the epineurium (EN, arrow) wrapping around outer surface of lamella of the OE (LOE). Lymphatic cells are found in OO (arrowheads) and OB. **C**. The LV extends centrally from the OO/OB along the ventral telencephalon (vTel) posteriorly to the ventral diencephalon. **D.** Higher magnification of OO in *B*. LV (arrowheads) is associated with olfactory nerve (ON, red) and covers ventral surface of OB (green, arrows). **E**. Lyve1b:EGF^+^ cells in tips of LOE resemble High Endothelial Venules (HEVs). **F**. Putative Mural lymphatic endothelial cells (MuLECs) wrap the OB (arrows). Representative images selected from detailed analysis of 9 brains. DAPI (blue). A, C**, =** 200 µm; B, D = 100 µm; C, F = 50 µm.

**Figure 2.**
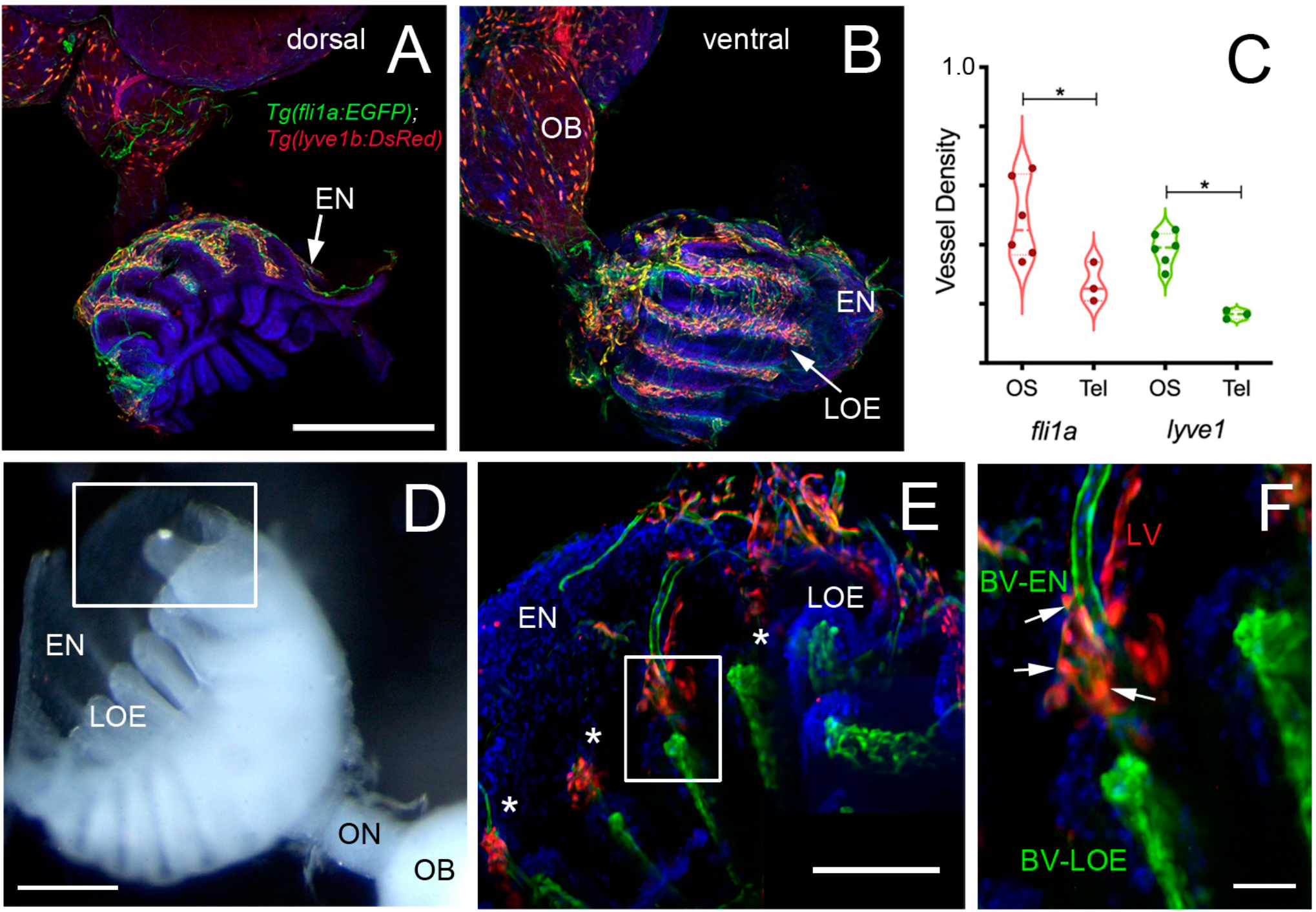
The adult olfactory organs (OO) have extensive and interconnected Blood (BV) and Lymphatic Vasculature (LV). **A, B**. Whole mount *Tg(fli1a:EGFP;lyve1b:DsRed)* adult OO connected to OB with BV (*fli1a:EGFP,* green) and LV *(lyve1b:DsRed*, red). Dorsal (A) and ventral (B) views; DAPI (blue), Scale Bars: A, B = 200 µm. **C**. BV (red) and LV (green) density is greater (SE, P-value <0.05, unpaired t-test) in olfactory system (OS = OE and OB) than telencephalon (Tel), *n* = 6 adult brains. One-way ANOVA, Tukey multiple comparison test, P < 0.05. Representative images selected from detailed analysis of least 6 brains. **D**. Transmitted light image of fixed whole mount OO. Boxed area represent where LOE connect with EN. Scale bar *=* 100 µm, **E**. At the distal tips of each lamellae (asterisks) the LV (red) meet the BV (green) (*E*, boxed area). Scale bar *=* 100 µm, **F**. Cells express both lyve1b:DsRed and fli1a:EGFP (arrows). Scale bar = 25 µm. A, B, E, F: Analysis of 9 brains.

**Figure 3.**
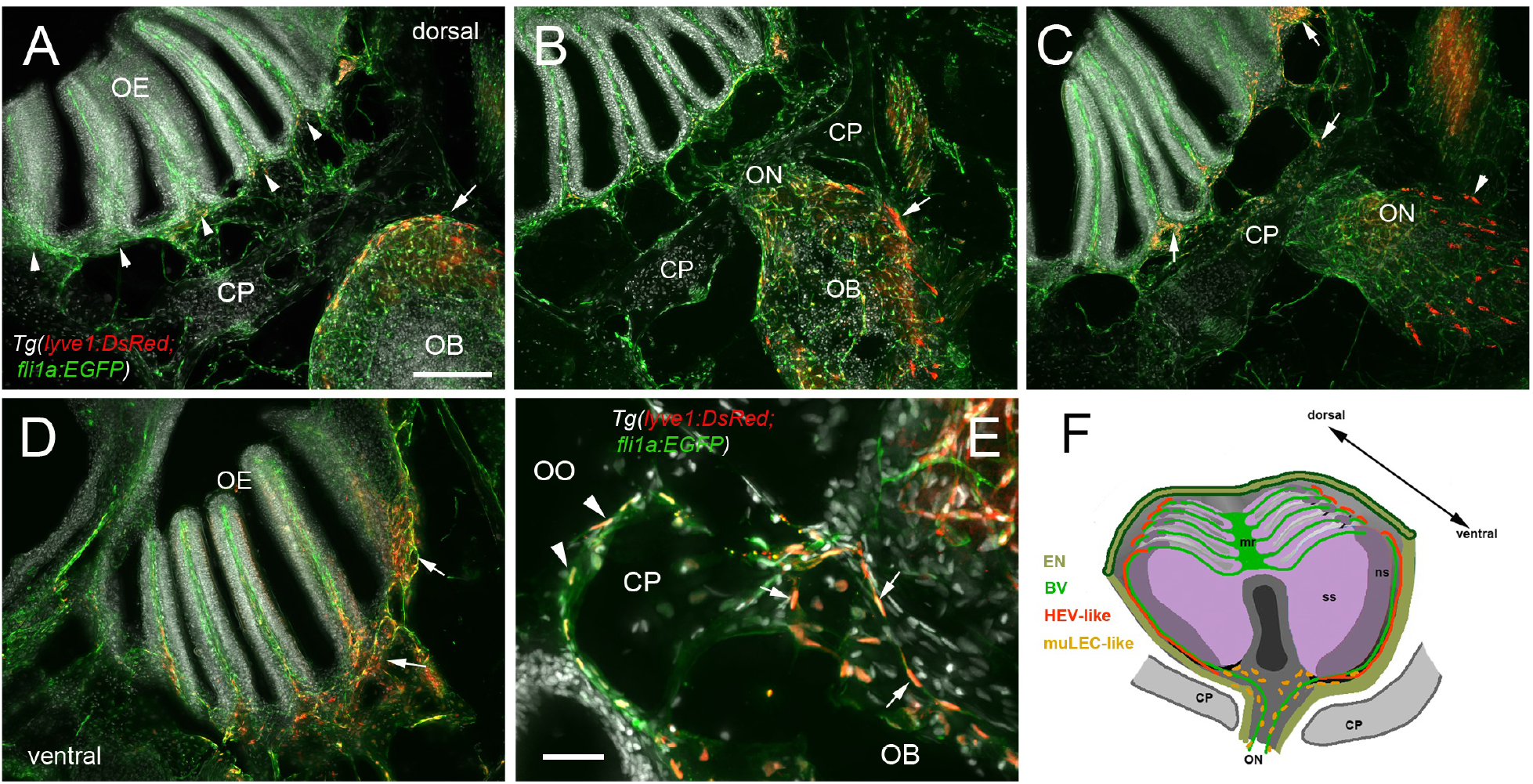
Blood vasculature extends through cribriform plate with muLEC-like lymphatic cells. **A-E.** Sections from *Tg(lyve1b:DsRed;fli1a:EGFP)* adult brains (n=6 brains). **A**. In dorsal sections the OB is separated from the OE by the cribriform plate (CP). The OB has extensive BV (green) extending into the lamellae of the OE and muLEC-like cells (red) on the surface of the OB (arrow). A-D = 100 µm **B**. The ON passes through the CP accompanied by extensive BV (green). muLEC-like cells are on the medial surface (red, arrow) of the OB. **C**. The muLEC-like cells (red, arrow) line the ventral side of the ON. **D**. muLEC-like cells line the basal OE (red, arrows) in the most ventral region of the OO. **E**. muLEC-like cells on the BV extending across the CP and many are positive for both lyvel1:DsRed and fli1a:EGFP (arrows). Scale bar = 50 µm. **F**. Diagram depicting olfactory organ with sensory (ss) and non-sensory (ns) epithelia that have extensive BV (green). The lamellae of the OE contain HEV-like LV (red) that do not extend across the cribriform plate (CP). muLEC-like cells (orange) line the BV and extend from the olfactory bulb across the CP to the basal OE. Scale bars: A-D = 100 µm, E = 50 µm.

To investigate the association of lymphatic vasculature (LV) with blood vasculature (BV) in the OOs, *Tg(lyve1b:DsRed;fli1a:EGFP)* animals were used to visualize the LV (red) and BV (green) (Fig. 2). We found extensive BV (fli1a:EGFP^+^) surrounding the OE associated with the EN in both the dorsal (Fig. 2A, green) and ventral (Fig 2B, green) OO and OB. The BV (Fig 2A, B, green) and LV (Fig. 2A, B, red) form an extensive network extending along the lamellae of the dorsal and ventral OE. In comparing the density of BV and LV in the dorsal brain, the OE/OB have a greater density of BV and LV than the telencephalon (Fig. 2C) reflecting the intimate association of the BV and LV with the OO/OB. The BV (Fig. 2A, B, E, F, green) and LV (Fig. 2A, B, E, F, red) extends along the EN that surrounds the LOE (Fig. 2D, E) and meet at the tips of the LOE where muLEC-L like cells were observed (Fig. 2E boxed area, red, F arrows). Thus the extensive BV and LV associated with the EN and OE connect along the distal lamellae where distinct BV morphologies are associated with the EN and LOE (Fig. 2F).

In mammals, the olfactory lymphatic route crosses the cribriform plate (CP) separating the OBs and OOs draining cerebral spinal fluid (CSF) through the perineural space surrounding olfactory nerve (Sun *et al*., 2018), connecting to nasal lymphatics and carrying lymphatic endothelial cells, T, B lymphocytes and antigen presenting cells (APCs) toward cervical lymph nodes (Kaminski *et al*., 2012). To characterize the LV structure crossing the cribriform plate we sectioned intact, decalcified heads from *Tg(lyve1b:DsRed;fli1a:EGFP)* animals to determine whether the muLEC-L cells or HEV-L cells extended across the cribriform plate (Fig. 3, CP). Dorsal to, and at the site of, ON crossing (Fig. 3, A, B) the OE was populated primarily by fli1a:EGFP^+^ BV. The lyve1b:DsRed^+^ LV (Fig. 3C, red, arrowhead) is associated with the fli1a:EGFP^+^ BV surrounding the ON (Fig. 3C, green) as it crosses the CP and lines the basal region of the OE (Fig. 3C, D, red, arrows). The muLEC-like cells of the LV lined the BV both on the intra-cranial (Fig. 3E, arrows) and extra-cranial side (Fig. 3E, arrowheads) of the ethmoid bone. We never observed HEV-L cells (Fig 1E) crossing the CP or on the intra-cranial side of the ethmoid bone. Thus the muLEC-L lymphatic cells associated with the BV were found wrapping the exterior surface of the OB (Fig. 1D, F), crossing the CP (Fig. 3) and extending along the EN (Fig. 3A, B) where they were associated with the HEV-L LV of the olfactory organ (Fig. 3F).

### Neutrophil populations in the adult olfactory organ

Neutrophils, the most abundant leukocyte sub-types in adult zebrafish, are essential players in the innate immune system and more recently have been shown to migrate not only on BV but also LV. We used the *Tg(OMP:RFP);Tg(mpx:GFP*) animals to visualize olfactory sensory neurons (red) and neutrophils (green), in fixed whole mount brains. Surprisingly, we observed neutrophils only in the OO of adult brains (Fig. 4A, B green). Neutrophils were localized in the fingerlike lamellae of the OE predominantly associated with the EN wrapping around the OE (Fig 2A, B). The OMP:RFP^+^ OSNs (Fig. 4A, B, red, ss, red) are in the central OE (ss) and peripheral regions of the lamellae are non-sensory epithelia (Fig. 4B, ns). The tips of the LOE are connected to the EN (Fig. 4B, EN, LOE, blue; Fig. 2). Analysis of the distribution of GFP-positive neutrophils revealed that they were located primarily in the ns epithelia and EN with many fewer neutrophils in the ss epithelia (Fig. 4B, E). Within the OE/EN there were three morphologically distinct mpx:GFP^+^ cells (Fig. 4C, D, F). Neutrophils with rounded shape (Fig. 4C, green, nt1) were associated with the basal OE, while neutrophils with amoeboid like morphology (Fig. 4C, green, nt2, D, ci=0.7) were present in the tips of the LOE and EN, although this distribution changed in response to damage of the OE (see below). In sectioned OE tissue the columnar shaped mpx:GFP^+^ cells (Fig. 4D1, green) were morphologically similar to sustentacular cells of the OE visualized with the *Tg(six4b:mCh)* reporter line ((Torres-Paz and Whitlock, 2014); Fig. 5A, B red). These cells lie at the interface of the ss and ns epithelia (Fig. 5C) and further studies are needed to carefully characterize this class of mpx:GFP^+^ cells. To confirm that the neutrophils observed in the whole mount OE (Fig. 4G, green) were within the OE as opposed to coating superficial layers, a z-stack analysis was performed (Fig. 4G, H) showing that the mpx:GFP^+^ cells are within the OE tissue. Thus the adult OOs are unique because they are the only regions of the adult brain where resident neutrophils are found under normal conditions.

**Figure 4.**
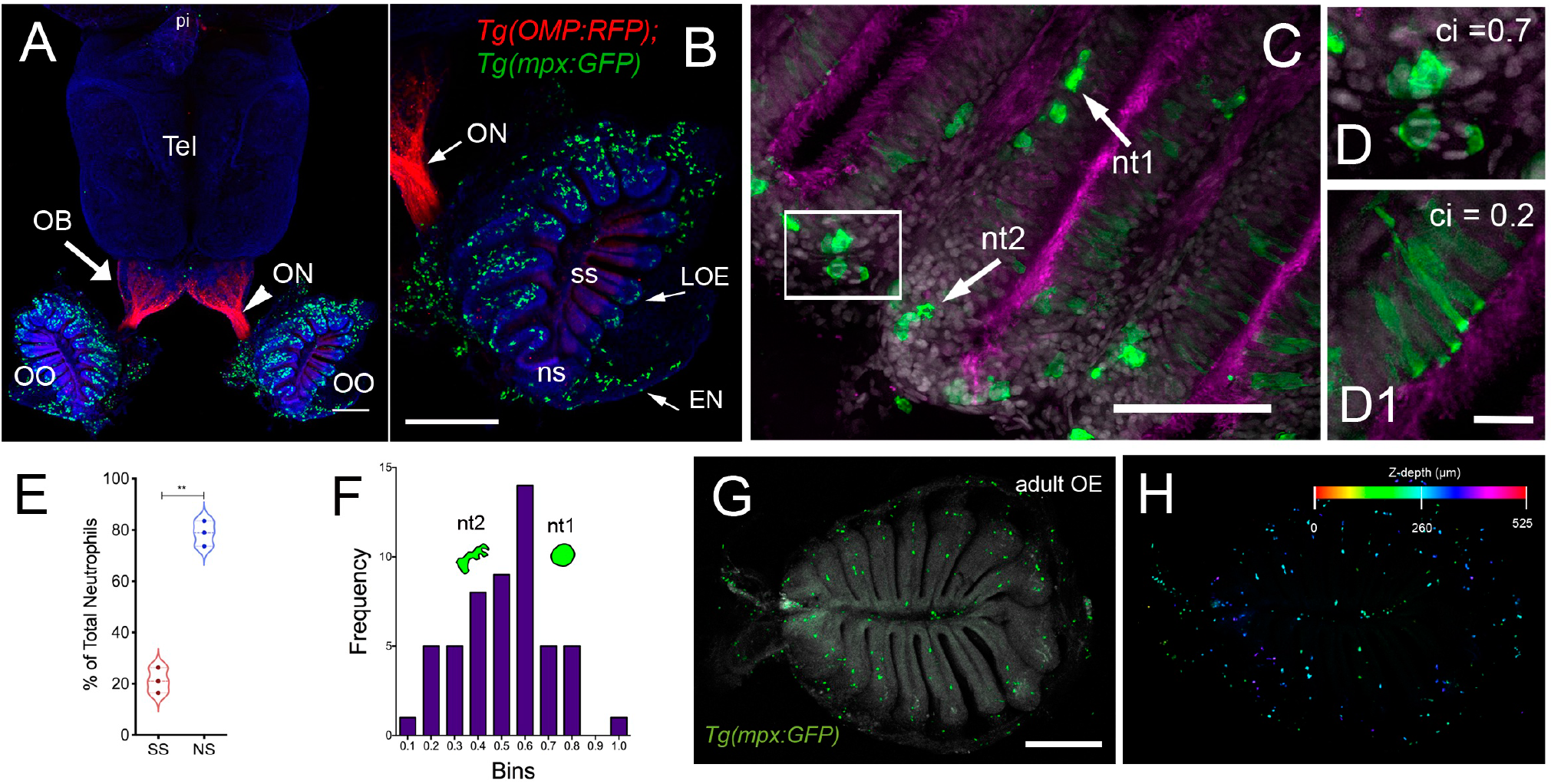
Neutrophils are found only in the olfactory organs of the adult brain. **A.** Wholemount brain of *Tg(OMP:RFP);Tg(mpx:GFP)* adult: neutrophils (green) are only present in the OO (OE/EN), telencephalon (tel), pineal (pi). Scale bars A, B = 200 µm. OO (from A) contains a large population of neutrophils (green, *n*=487 neutrophils). OMP:RFP^+^ OSNs are located only in sensory epithelia (ss, red) not in non-sensory epithelia (ns), olfactory nerve (ON). **C**. Neutrophils with a rounded shape, (nt1, arrow: circularity index 0.7 or greater) and amoeboid shape (nt2, arrow: circularity index of 0.4-0.6; **F**) were observed in the LOE. **D**. Neutrophils with an amoeboid shape (nt2, arrow) were located throughout the OE and EN. **D1**. Sustentacular-like cells (**D1**), circularity index 0.2, lie at ns-ss epithelia interface (Fig. 5). **E**. Total number of neutrophils in the OO. The non-sensory (ns) tissues (respiratory epithelia + NE, blue) have more neutrophils than sensory epithelia (ss, red), n= 3 OE from 3 different fish. **F**. Frequency distribution of nt1 and nt2 cells (*n*= 53 neutrophils from brain shown in C). **G**. Maximal projection of whole mount *Tg(mpx:GFP)* adult OE: Neutrophils (green); autofluorescence (gray). (*H*) Neutrophils (from *G*) were color-coded based on (H) Z-stack depth. Total depth= 550µm. Scale bars A, B = 200 µm; C = 60 µm; D, D1=20 µm; G, H =100 µm. A, B: 9 brains imaged; C, D: 6 brains sectioned

**Figure 5.**
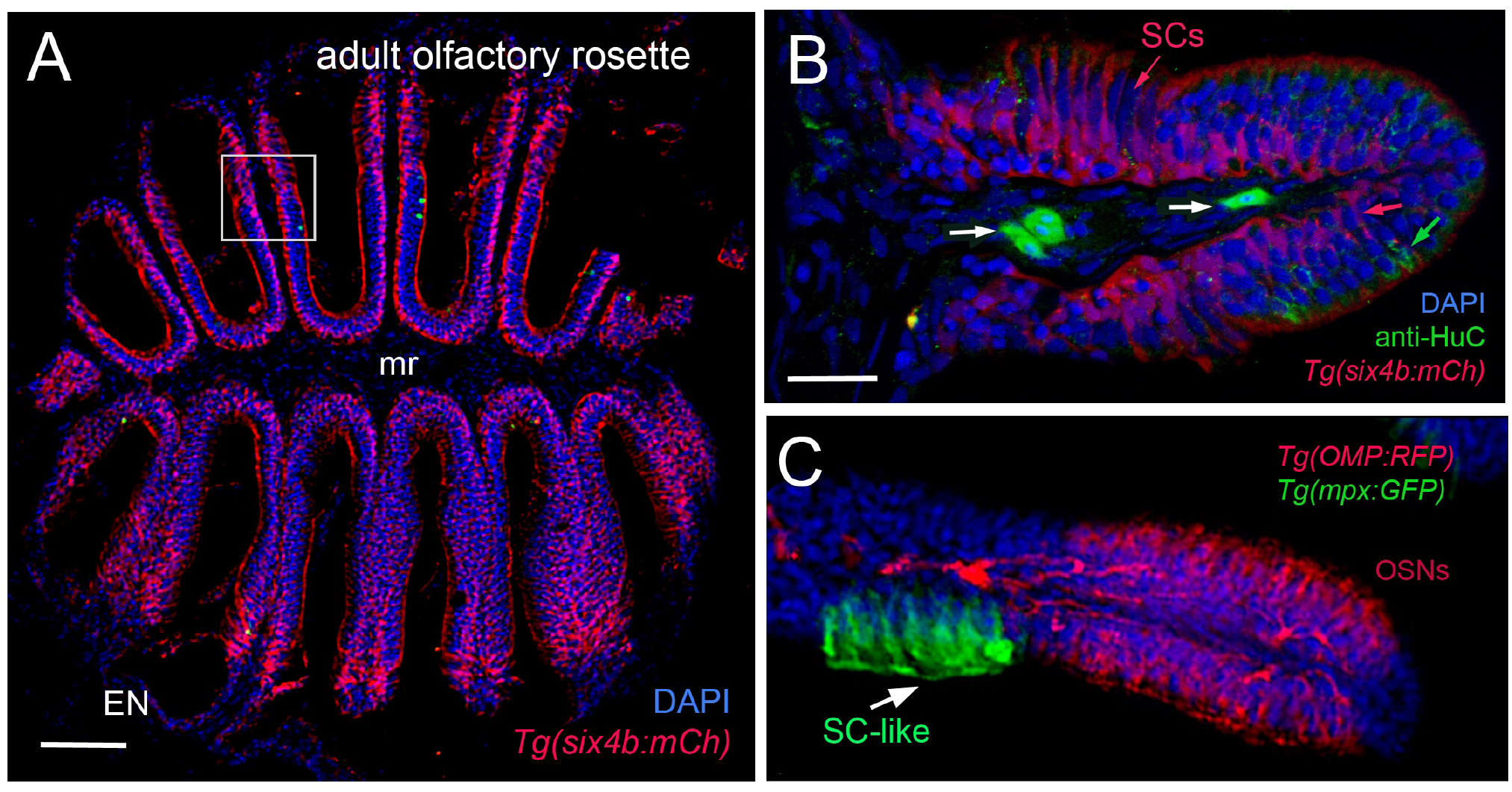
Sustentacular cells in the OE are associated with markers for neutrophils. **A-C.** Cryosections of adult olfactory epithelia. **A**. Low magnification of adult olfactory rosette (OE) from *Tg(six4b:mCh)* line showing sustentacular cells (red) that are distributed within the lamellae of the OE where some areas have denser clusters (boxed area). Epineurium (EN), midline raphe (mr). Scale bar = 100 µm. **B**. Lamellae of OE with Six4b:mCh^+^ SCs (red) and anti-HuC^+^ neurons (green). Scale bar B, C = 25 µm.**C.** mpx:GFP^+^ cells (green, arrow) lie in clusters adjacent to OSN (red) and are similar to SCs (SC-like, see *B*, red). A, B: 1 sectioned brain, C: 6 sectioned brains.

### Neutrophil response to damage in the adult olfactory sensory system

In order to investigate the neutrophil response to damage of the OE, we exposed *Tg(mpx:GFP);Tg(OMP:RFP*) adult fish to 10 µM CuSO4. Because of the challenges of live imaging in the whole mount adult brain, adults were sacrificed at different times after copper exposure to follow the dynamics of neutrophil response over time. In untreated control animals, consistent with previous results, neutrophils (Fig. 6, green) were observed only in the OO (Fig. 6A, arrowhead, A’) and were absent in the brain (Fig. 6A). After four hours of copper exposure, an increase in neutrophils was observed in the OO (Fig. 6B, green, arrowhead, B’, B’’). Within the OO the ns and ss OE as well as the EN (Supp. Fig. 1F, H) showed an increase in neutrophils in response to damage. Additionally neutrophils were observed in the ventromedial OB, along the telencephalic ventricle (Fig. 6B, OB, V) and in the ventral telencephalon (Fig. 6B, green, arrows). Fish left to recover for one day post-treatment still showed elevated numbers of neutrophils in the ventral OB (Fig. 6C, green, arrows, *D*) and the OO (Fig. 6C’, green). The increased numbers of neutrophils in the OOs and subsequent appearance of neutrophils in the ventral OB and ventral telencephalon (Fig. 6D, E, vCNS), suggests that neutrophils may move from the OOs into the ventral CNS in response to peripheral damage.

**Figure 6.**
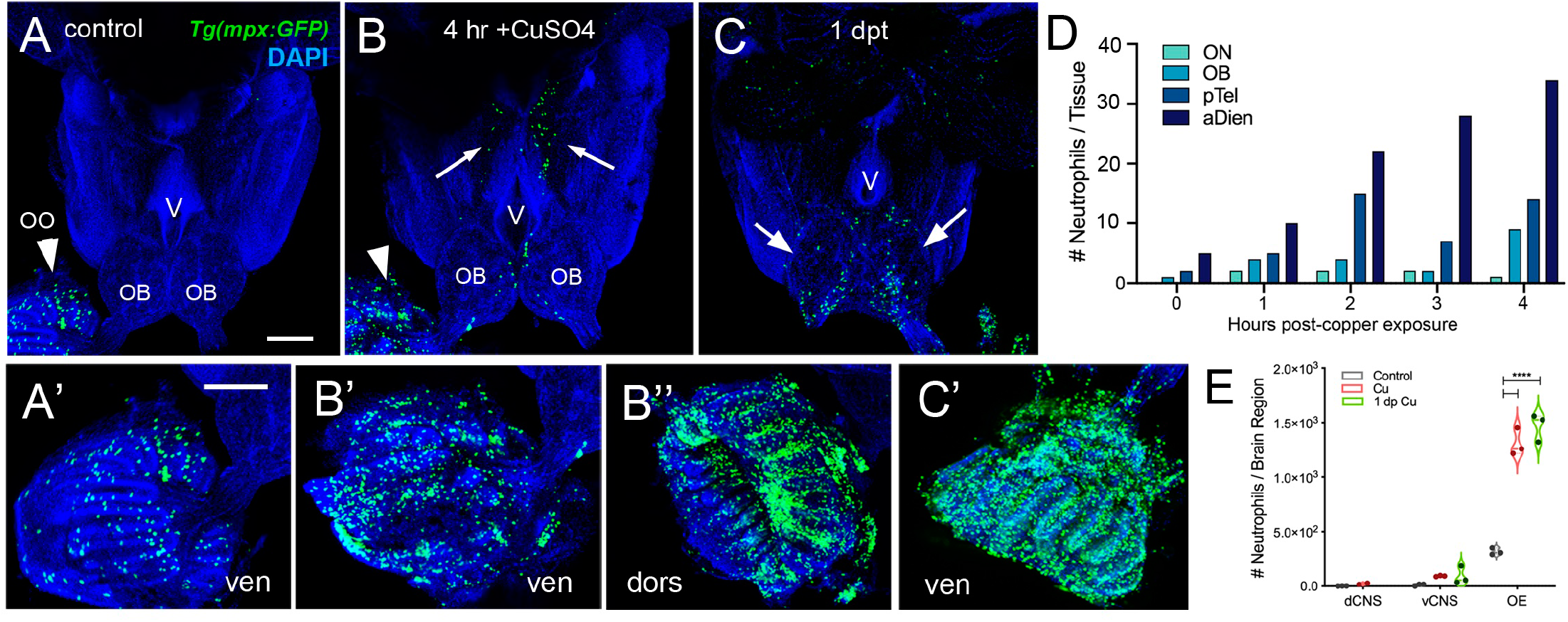
Exposure to copper is correlated with increased neutrophils in the peripheral and central nervous system. **A-C.** Ventral views of whole mount adult brains from *Tg(mpx:GFP)*. Scale bars: A-C; A’-C’= 100 µm. **A**, **A’**. Control with neutrophils found only in OO (arrowhead; A’). (**B.** After four-hour exposure to copper, there is an increase in the number of neutrophils in the OO (B, arrowhead, B’, B”). Neutrophils were observed in the ventral OB, along the ventricle (V) and in the ventral telencephalon (B, arrows). **C**. One day post treatment neutrophils are still present in OO (C’), OB (arrows) and ventral telencephalon. **D**. Neutrophils appear over time in an anterior to posterior spatial pattern in the CNS. OO is not plotted because number (average ∼1,500) is out of range (see E). **E**. Copper exposure was correlated with increased neutrophils in OE and ventral CNS. A-C, A’-C’: 3 brains were examined per treatment and summarized in E. Preparations were selected for imaging based on whether they were intact and the signal to noise of the labeling. D:For each timepoint 1 brain was analyzed.

### Damage induced changes in cell cycle dynamics in the olfactory sensory system

To further investigate the cellular dynamics of the neutrophil response to copper-induced damage in the adult, we repeated the experiments with copper using *Tg(mpx:GFP)* animals in the presence of BrdU. When viewed in flattened whole mount preparations (Fig. 7A-C), the OE of the adult is organized as a “rosette” with the central region midline raphe (mr) surrounded by ss and the outer regions of the rosette (tips of the lamellae) containing the ns or respiratory epithelia. In control animals (Fig. 7A, viewed looking into the rosette) BrdU labeling, consistent with the mitogenic nature of the olfactory system, was observed (Bayramli *et al*., 2017; Brann and Firestein, 2014). After four hours of copper exposure, BrdU labeling showed significant increases in the mr (Fig. 7B, white, arrow), and in the ns epithelia extending to the EN. In contrast, one day post treatment (dpt) significant increases in BrdU labeling were observed in the ss epithelia (Fig. 7C, F) consistent with the renewal of OSN in the OE after damage (Iqbal and Byrd-Jacobs, 2010). Additionally the neutrophils now lined LOE (Fig. 7C, green) possibly in association with the BV (Fig. 7D, green). The number of neutrophils showed significant increases at 4 hours post-treatment (hpt) and remained high in the ss epithelia one dpt (Fig, 7E; 444.67 ± 31.39 and 373.33 ± 32.32 neutrophils in 4 hpt, red, and 1 dpt, green). Significant increases in BrdU labeling at both 4 hpt and 1 dpt were observed only in the ss epithelia (Fig. 7F; 480 ± 241.76 and 786 ± 211.6, respectively). Analysis of cells expressing both mpx:GFP and BrdU showed a significant increase compared to control animals (Fig. 7G, control: 9 ± 1, 4 hpt: 26 ± 6, 1 dpt: 22 ± 5.29). The frequency of rounded neutrophils (see Fig. 4C, green, nt1; ci 0.7 or greater) and amoeboid-like (see Fig. 4C, green, nt2; ci 0.4-0.6), potentially representing “resting” and activated neutrophils, respectively, increased in the OE post-damage (Fig. 7H). The columnar shaped cells (ci 0.1-0.3) increased in frequency at one dpt in the sensory region (Fig. 7H, green, 0.2 -0.3 green bars) but remained as the least common morphology. We found that damage to the OE resulted in an increased number of rounded neutrophils and a small but significant number were double labeled for BrdU, thus the majority of the increase in neutrophil number was likely due to migration as opposed to proliferation. Future work using photoconvertible lineage tracers will allow us to determine the exact contribution of local vs. immigrant neutrophils in the response to damage in developing and adult brain.

**Figure 7.**
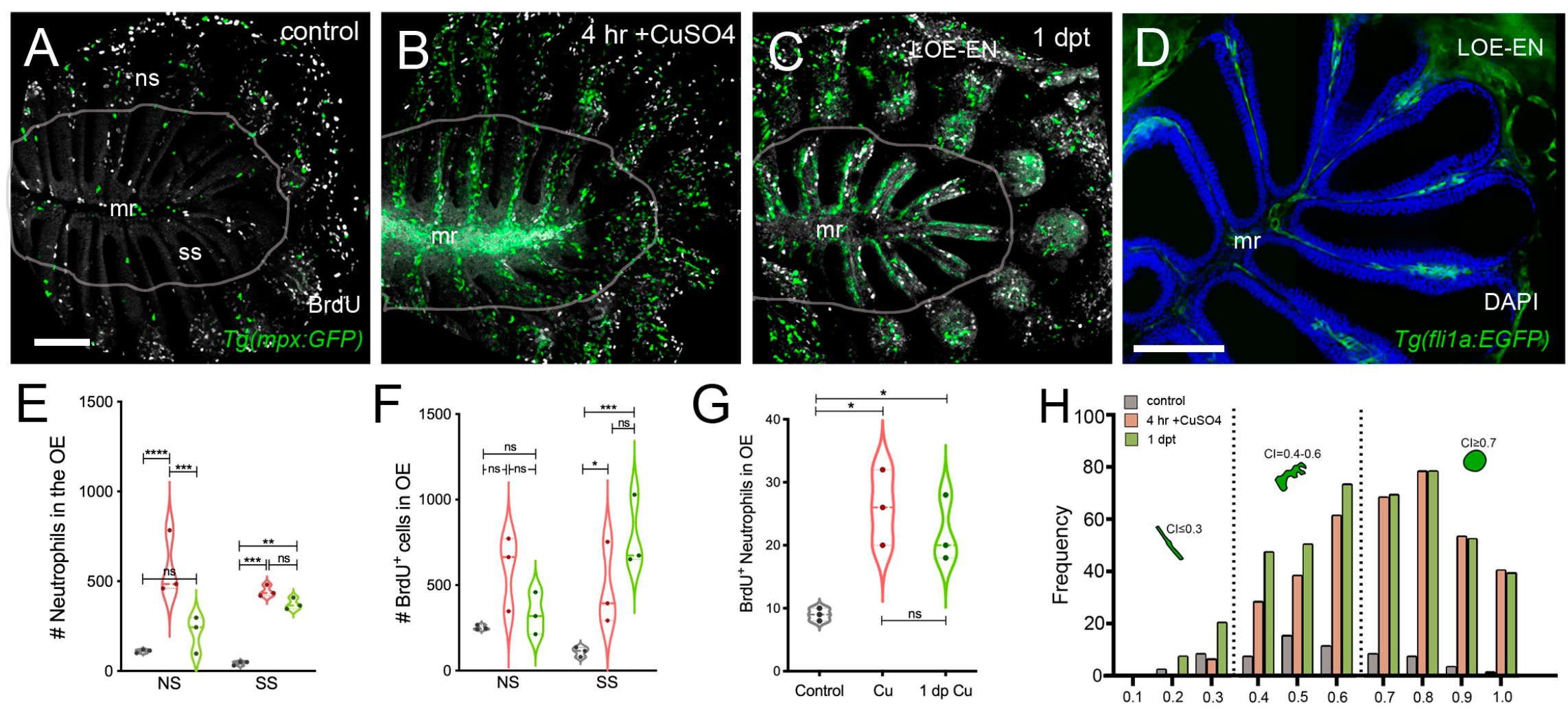
Damage induces changes in cell division of OSN and neutrophil precursors in the adult olfactory organ. **A-C**. BrdU labeled cells (white), neutrophils (green) in whole mount OO of adult fish. Scale bars A-C = 50 µm **A**. Prior to copper exposure BrdU labeling and scattered neutrophils were observed in the medial raphe (mr), sensory (ss), and non-sensory (ns) epithelia. n= 3 OE. **B**. After four hours of exposure to copper intense BrdU labeling was observed in the mr. n= 3 OE. **C**. One day post recovery neutrophils lined the lamellae and intense BrdU labeling was observed in ss and LOE-EN. n= 3 OE. **D**. Section of *Tg(fli1a:EGFP)* adult OE showing extensions of blood vasculature (green) within the OE. Scale bar = 100 µm. n= 3 sectioned heads. **E.** Significant increases in neutrophil number were observed after 4 hour copper exposure (red) in both the ns and ss epithelia when compared to control (grey). At one dpt (green) only the ss remained significantly greater than controls. n= 3 OE. **F**. Damage induced changes in BrdU^+^ cells have were significant in the ss epithelia but not the ns at 4 hour copper exposure (red) and one dpt (green). n= 3 OE. **G**. There was a small but significant increase in mpx:GFP^+^ cells double labeled for BrdU scored in the OE. (E-G, *n*=3 adult OE from different fish; Two-way ANOVA, Tukey multiple comparison test, p < 0.05). (*P, 0.05, **P, 0.01, ***P, 0.001). n= 3 OE. **H**. Four hours of copper exposure (orange) and one day post-treatment (green) resulted in an increase in rounded neutrophils (nt1; circularity index 0.7 or greater see Fig. 1) and amoeboid neutrophils (nt2; circularity index 0.4-0.6) when compared to controls.

## DISCUSSION

In this study we have shown that the olfactory sensory system has a unique “immune architecture” where neutrophils permanently populate the olfactory sensory organs in association with a complex network of BV-LV. These neutrophils mount a rapid response to copper-induced damage to the OE populating not only the tissues of the OE and associated EN, but also appearing in tracts extending posteriorly along the ventral CNS. These data demonstrate a role for resident neutrophils in the olfactory sensory system and suggest that the nasal lymphatic pathway may be a potential site of entry for immune cells into the CNS.

### Lymphatic Vasculature

The olfactory/nasal lymphatic route was first described using India ink to label CSF drainage pathways from the brain where particles moved from cranial subarachnoid space to lymphatic channels of the olfactory mucosa (Jackson *et al*., 1979). Subsequently it was shown that while the subarachnoid space of the optic nerves and cochlea region were labeled, the only direct connection between cranial CSF and lymphatics was the nasal route (Kida *et al*., 1993; Kida *et al*., 1995; Koh *et al*., 2005) passing through cribriform plate along perineural spaces near the olfactory nerves to the nasal mucosa and cervical lymph nodes (Sun *et al*., 2018). With the re-discovery of the brain lymphatics (Louveau *et al*., 2015) the relative importance of the drainage of CSF via the meningeal LV versus olfactory/nasal LV is currently a subject of debate (see for discussion (Dolgin, 2020).

In descriptions of the olfactory/nasal drainage in mammals, the LV is generally depicted with terminations at the extra-cranial side of the cribriform plate. Here we found two types of lyve1b:EGFP^+^ LV: one having muLEC like structure where the cells line the BV (Bower and Hogan, 2018; Bower *et al*., 2017), appeared to be connected, and were found on both the intracranial and extra cranial side of the cribriform plate; and a second with morphology similar to HEVs that were found in association with the OE/EN on the extra-cranial side of the cribriform plate.

The muLEC-like LV wrap the dorsal and ventral surfaces of the olfactory bulbs extending posteriorly along the ventral telencephalon and anteriorly through the cribriform plate with the BV. Within the muLEC-like cells there were two populations: one positive only for lyvel1b and a second positive for both lyve1b and fli1a. During development muLECS have been shown to form from local blood vessels by (Bower *et al*., 2017) and these forming cells are positive for both fli1a and lyve1b. Thus, the lyve1b+/fli1a+ population may represent adult progenitors of LV important in restructuring the OE after extensive damage. The muLEC-like cells appear to be connected, yet future studies are needed to confirm that these cells are from the non-lumenized mural lineage (Okuda and Hogan, 2020).

#### Lymph node equivalent in fish

In mammals the nasal lymphatic route that drains into the cervical lymph nodes through the cribriform plate, carry immune cells such as monocytes, dendritic cells, and T cells (Goldmann *et al*., 2006; Hsu *et al*., 2019). In addition, mammals have Nasal-Associated Lymphoid Tissue (NALT) also referred to as Waldeyer’s lymphatic ring, surrounding the naso/oropharynx. This tissue contains lymphatic vessels and HEVs, which are specialized post-capillary venous swellings, enable lymphocytes circulating in the blood to directly enter a lymph node (by crossing through the HEV). Recently tissue described as NALT has been reported in fish (Das and Salinas, 2020; Sepahi and Salinas, 2016), yet fish do not have lymph nodes. Thus a distinction is made between “organized” NALT and “diffuse NALT” (Sepahi and Salinas, 2016) or NALT versus non-NALT (for murine nasal dendritic cells (Lee *et al*., 2015) where teleost fish have diffuse-NALT/non-NALT in the olfactory organs. Here we found that the OE/EN has an extensive blood vasculature associated with lyve1b:EGFP^+^ lymphatic endothelial cells resembling high endothelial venules (HEVs) of the lymph nodes in mammals (Fig. 8, olfactory epithelia, upper, HEV-like). The HEV-like cells were localized to the tips of the LOE extending on the external side of the EN to the base, terminating in the region where the meningeal membranes fuse on the extra-cranial side of the cribriform plate. At the tips of the LOE, on the internal side, the BV is associated with the HEV-like cells and in this region we identified lyve1b/fli1a^+^ cells similar to those seen in the OB although not on the BV. This cell type was also observed lining the cribriform plate in the region of the meninges (Fig. 5). The structures observed raise the possibility that in spite of lacking lymph nodes, the zebrafish OOs shows similarities with mammalian lymph node organization thus suggesting the existence of an organized secondary lymphoid tissue in the OO.

**Figure 8.**
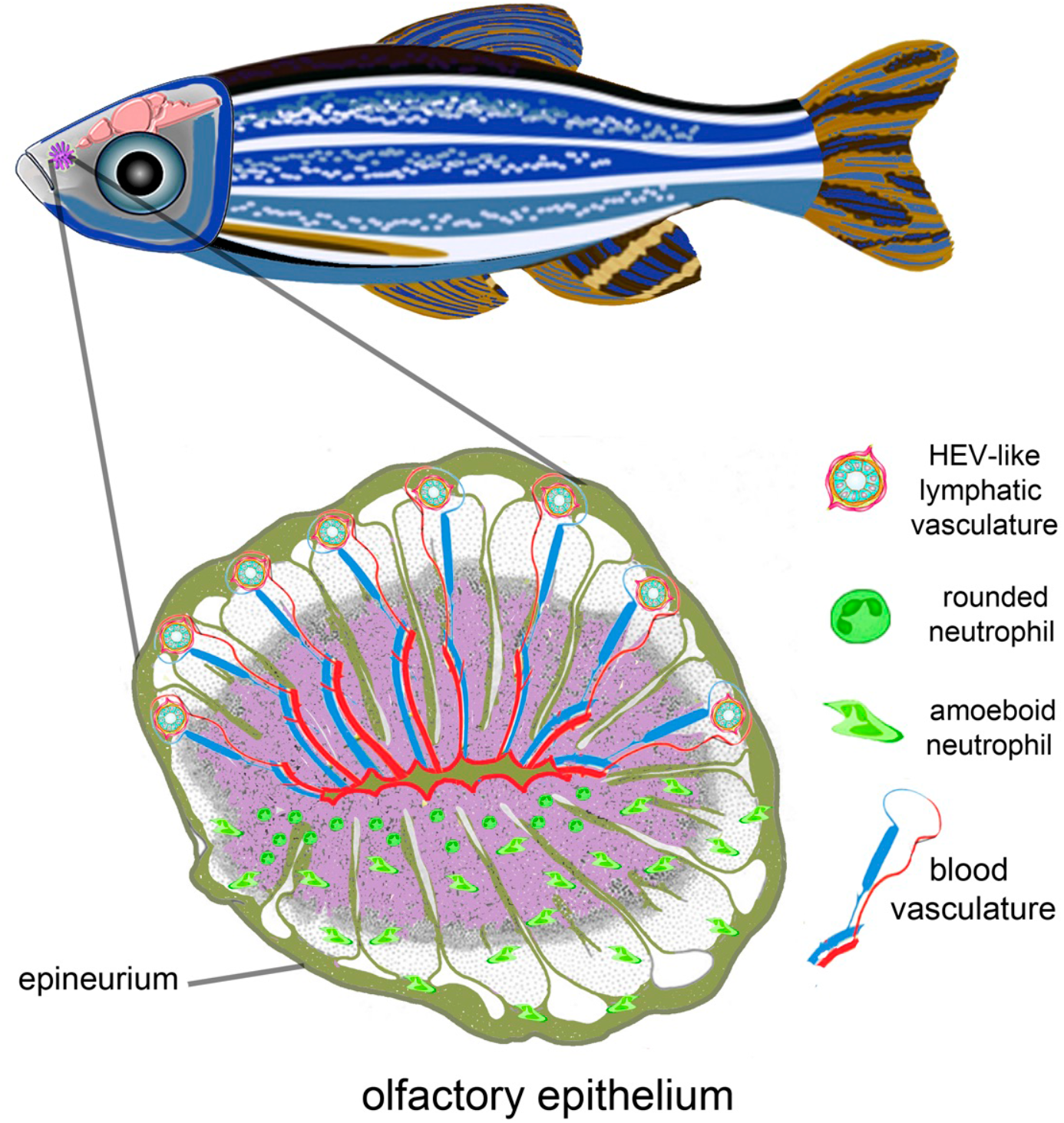
Olfactory organ is neural-immune interface. Schematic of olfactory epithelium of adult zebrafish. The Blood-Vasculature-LV and neutrophils are shown in different halves of the olfactory rosette for clarity. Upper half of olfactory epithelium are proposed connections of Blood-Vasculature (BV) with LV via HEV-like cells that may also contact the meningeal immune system via vasculature associated with the epineurium. Lower half of olfactory rosette depicts resident neutrophils found only in the OE.

#### Neutrophils

It has recently been shown that neutrophils, in addition to their role as the first line of defense in the innate immune response, also transport antigens and populate lymph nodes via HEVs where they coordinate early adaptive immune responses (Hampton *et al*., 2015; Hampton and Chtanova, 2016; Li *et al*., 2019). Neutrophils are found in many tissues and these subpopulations of neutrophils perform many functions (Rosales, 2018) such as in the lung, which is known to retain neutrophils as a host defense niche (Kubes, 2018; Yipp *et al*., 2017). In mammals the OE is reported to have B lymphocytes, lactoferrin and lysozyme, in the Bowman’s glands (Mellert *et al*., 1992) and neutrophils in the non-sensory epithelium of the vomeronasal organ (Getchell and Kulkarni, 1995). In teleosts, limited morphological studies have shown scattered myeloid and lymphoid cells within the OE and lamina propria (Dong *et al*., 2020; Tacchi *et al*., 2014) (Yu *et al*., 2018). Most recently, in the OO, in response to inflammation neutrophils infiltrate and later express neurogenesis-related genes suggesting a potential role for neutrophils in the ongoing neurogenesis of the OE (Ogawa *et al*., 2021).

The neutrophils we observed in the OOs were striking not only in their number but also their limited distribution: they were found only in the OOs of the adult brain under normal conditions. After copper exposure there was a large increase in the number of neutrophils in the OE/EN and subsequently neutrophils appeared in the CNS initially in the ON, ventral lateral OB and then extending posteriorly along the ventral telencephalon, although far fewer neutrophils were observed in the CNS. This ventral tract from OOs contains a rich network of LV (Fig. 1), and has previously been suggested as a route for immune cell influx through the basal forebrain in mice (Pägelow *et al*., 2018) and mesenchymal stem cell migration cell from the periphery to the OB (Galeano *et al*., 2018). Thus the pattern of neutrophils observed is suggestive of neutrophil movement from the periphery along the ON, ventral OB and ventral telencephalon. While it is tempting to propose that these neutrophils enter from the OE into the CNS, more experiments are needed to better understand the source of the CNS neutrophils.

#### Conclusions

In mammals the olfactory/nasal brain lymphatic drainage system is assumed to function not only in water homeostasis and pressure regulation, but also in immune responses and surveillance within the meningeal lymphatic system (Sun *et al*., 2018). Yet here we have shown that the OOs have an extensive blood lymphatic vasculature (including HEV-like structures) enveloping the OE, a large resident neutrophil population and furthermore, that damage induced in the olfactory sensory epithelia is correlated with the appearance of neutrophils in the brain. Whether the presence of these neutrophils is related to the regenerative properties of the OE as the OSNs undergo constant replacement, represents a special population secondary lymphoid tissue capable of mounting a rapid immune response, or both, remains to be determined.

## Material and Methods

### Animals

Zebrafish were maintained in a re-circulating system (Aquatic Habitats Inc, Apopka, FL) at 28°C on a light-dark cycle of 14 and 10 hours respectively. All fish were maintained in the Whitlock Fish Facility at the Universidad de Valparaiso. Wild-type (WT) fish of the Cornell strain (derived from Oregon AB) were used. All protocols and procedures employed were reviewed and approved by the Institutional Committee of Bioethics for Research with Experimental Animals, University of Valparaiso (#BA084-2016). Adults used in the study were 12-16 months of age. The following transgenic lines were used to visualize specific cell types: *Tg(BACmpx:gfp)^i114^, Tg(mpx:GFP) (Renshaw et al., 2006)*; (*Tg(fli1a:EGFP)^y1^ Tg(fli1a:EGFP*; (Lawson and Weinstein, 2002); *Tg(−5.2lyve1b:DsRed)^nz101^, Tg(2lyve1b:DsRed) Tg(−5.2lyve1b:EGFP)^nz151^ Tg*(*lyve1b:EGFP*), (Okuda *et al*., 2012); *Tg(gata1a:DsRed)^sd2^ Tg*(*gata1a:DsRed*) (Traver *et al*., 2003), *Tg(pOMP^2k^:gap-YFP)^rw032a^*, *Tg(OMP:YFP); Tg(pOMP^2k^:lyn-mRFP)^rw035a^ Tg(OMP:RFP*) (Sato *et al*., 2005); *Tg(six4b:mCh)*, (Harden *et al*., 2012).

### Copper Exposure

Initial dose response analysis was performed based on previous work in zebrafish and salmon (Baldwin *et al*., 2003); (Hernandez *et al*., 2011). A stock solution of 10 mM CuSO_4_ was diluted in system water for a final concentration of 10 uM CuSO_4_.

### Immunocytochemistry and Cell Labeling

Dissected adult brains were fixed in 4% PFA in 0.1M phosphate buffer 0.4M pH 7.3), or 1X phosphate-buffered saline PBS pH 7.4. Brains were rinsed three times in phosphate buffer or PBS, permeabilized in acetone at −20 °C for 10 minutes and then incubated for two hours in blocking solution (10 mg/ml BSA, 1% DMSO, 0.5% Triton X-100 (Sigma) and 4% normal goat serum in 0.1M phosphate buffer or 1X PBS). Primary antibodies used were anti-RFP (rabbit 1:250, Life Technologies), anti-GFP (mouse 1:500, Life Technologies), anti-GFP (rabbit 1:500, Invitrogen), anti-DsRed (mouse 1:500, Santa Cruz Biotechnology), anti-HuC/D (rabbit 1:500, Invitrogen) and anti-BrdU (rabbit 1:250,Invitrogen). Adult brains were incubated with the primary antibody for up to a week. After washes, tissues were incubated overnight in experiment dependent secondary antibodies: Dylight 488 conjugated anti-mouse antibody (goat 1:500, Jackson Immuno Research), Alexa Fluor 488 conjugated anti-rabbit antibody (goat 1:1000, Molecular Probes), Alexa Fluor 568 conjugated anti-rabbit antibody (goat 1:1000, Molecular Probes), Alexa Fluor 568 conjugated anti-mouse antibody (goat 1:1000, Molecular Probes), Dylight 650 conjugated anti-rabbit antibody (goat 1:500, Jackson Immuno Research), Alexa Fluor 350 conjugated anti-rabbit antibody (goat 1:1000, Molecular Probes). Tissues were then rinsed in 0.1M phosphate buffer or 1X PBS with 1% DMSO, stained for DAPI (1 µg/ml, Sigma), washed in 0.1M phosphate buffer or 1X PBS and mounted in 1.5 % low melting temperature agarose (Sigma) in an Attofluor Chamber for subsequent imaging (see below).

### BrdU Labeling

For each experiment nine adult fish were first housed overnight in 1.5 liter tanks containing 10 mM BrdU in system water. The next morning three fish were transferred to a new 1.5-liter tank with system water (control) and six fish were transferred to a new 1.5 liter tank with system water containing 10 μM CuSO_4_, and allowed to swim freely (4 hours). All control fish (3) and half of copper-exposed fish (3) were then anesthetized, sacrificed and heads fixed overnight in 4% PFA/1X PBS. The other half of copper-exposed fish (3) were transferred to a clean 1.5-liter tank, filled with system water, and allowed to recover. The next day, these fish were anesthetized, sacrificed and fixed as described above. After fixation, heads were incubated in EDTA (0.2 M, pH 7.5) for three days at 4 °C and brains dissected in sterile 1X PBS and pre treated in 2 M HCl for 30 minutes at 37 °C. Immunocytochemistry was performed as described in Immunocytochemistry & Cell Labeling. For imaging, whole adult brains were mounted on 2% low melting temperature Agarose, and OE were mounted between coverslips, as described above. The removal of brains from the skull with the OO still attached is a difficult dissection because the OSN axons pass through the cribriform plate to arrive in the OB. Therefore it was not always possible to have a preparation with both OE still connected to the brain.

### Cryosectioning

Fish were euthanized and heads were fixed overnight in 4% PFA at 4 °C and decalcified in EDTA (0.2 M, pH 7.6) for 3 days, and later embedded in 1.5% agarose/ 5% sucrose blocks and submerged in 30% sucrose for 3 days at 4 °C. Blocks were frozen (−20 °C) with O.C.T. Compound (Tissue Tek®) and sectioned (25 µm) using a cryostat. For flat mounting of the olfactory epithelia, olfactory rosettes were dissected after immunohistochemistry or staining, and mounted with the caudal side down on Poly-L-Lysine coated slides between triple 22×22 coverslip bridges and covered in VECTASHIELD® Antifade Mounting Media (Vector laboratories).

### Imaging and Image analysis

*Microscopy:* Fluorescent images were taken using a Spinning Disc microscope Olympus BX-DSU (Olympus Corporation, Shinjuku-ku, Tokyo, Japan) and acquired with ORCA IR2 Hamamatsu camera (Hamamatsu Photonics, Higashi-ku, Hamamatsu City, Japan). Images were acquired using the Olympus CellR software (Olympus Soft Imaging Solutions, Munich, Germany). Some images were also obtained using a confocal laser scanning microscope (Nikon C1 Plus; Nikon, Tokyo, Japan). Images were then deconvoluted in AutoQuantX 2.2.2 (Media Cybernetics, Bethesda, MD, USA) and processed using FIJI (National Institute of Health, Bethesda, Maryland, USA; (Schindelin *et al*., 2012) and CellProfiler (McQuin *et al*., 2018).

### Image Analyses

*Neutrophils*: Only neutrophils within the boundaries of the olfactory organs in adults were counted and the values were given as the average of total number of mpx:GFP positive with standard deviation. Values given for paired sensory structure are a sum of the individual sensory tissues.

To analyze the distribution of mpx:GFP^+^ neutrophils from both whole adult brains and flat-mounted olfactory rosettes, images were filtered by size (6-30 µm) and pixel intensity, and then counted using CellProfiler available Pipelines (McQuin *et al*., 2018). For quantification of neutrophils in different regions of the OE, sensory (ss) versus non-sensory (ns) regions were separated using *Tg(OMP:RFP)* animals or anti-HuC/D labeling as neuronal markers. We grouped the ns region with the epineurial extensions (EN) wrapping the OE. The percent of total neutrophils is the number of GFP cells in ss or ns regions, divided by total (sum of all GFP positive cells in ss, ns and EN). BrdU nuclei were detected by filtering size between 2-5 µm and co-localization between BrdU and neutrophils was done using “Co-localization” Pipeline in CellProfiler (McQuin *et al*., 2018).

The circularity index of each neutrophil was calculated using Analyze Particles in FIJI (National Institute of Health, Bethesda, Maryland, USA; (Schindelin *et al*., 2012). Neutrophils were size-filtered and values were graphed according frequency of distribution.

#### BV/LV vessel density

Density is defined by the ratio of the area positive for fli1a:EGFP (BV) and lyve1b:DsRed (LV) over the total dorsal telencephalic or the olfactory system area (which includes both the OE and OB). Protocol adapted from Zhao et al., 2016 (Zhao *et al*., 2016).

#### Statistics

Data are presented as means ± standard deviations. Experiments number and statistical analysis were done using Prism 9 (Graphpad), and are indicated in each figure legend. Unpaired Student’s t-tests were performed unless otherwise indicated. P values are indicated as follows: *P, 0.05, **P, 0.01, ***P, 0.001.

## Acknowledgments

We would like to acknowledge Andrea Moscoso and Maria Trinidad Ordenes for excellent management of the zebrafish facility.

## Funding

Grants/Fellowships Fondo Nacional de Desarrollo Científico y Tecnológico (FONDECYT) 1160076 (KEW); ICM-ANID Instituto Milenio Centro Interdisciplinario de Neurociencias de Valparaíso PO9-022-F, supported by the Millennium Scientific Initiative of the Ministerio de Ciencia (K.E.W, M.F.P.); CONICYT Doctoral Fellowship (ANID) 21161437 (MFP). The funding bodies did not take part in the design of the study, the collection, analysis, and interpretation of data, or in the writing of the manuscript.

## Availability of data and materials

The datasets used and/or analyzed during the current study are provided as a supplemental Source Data file.

## Competing Interests

The authors have no competing interests.

## SUPPLEMENTAL FIGURES

**Supplemental Figure 1.**
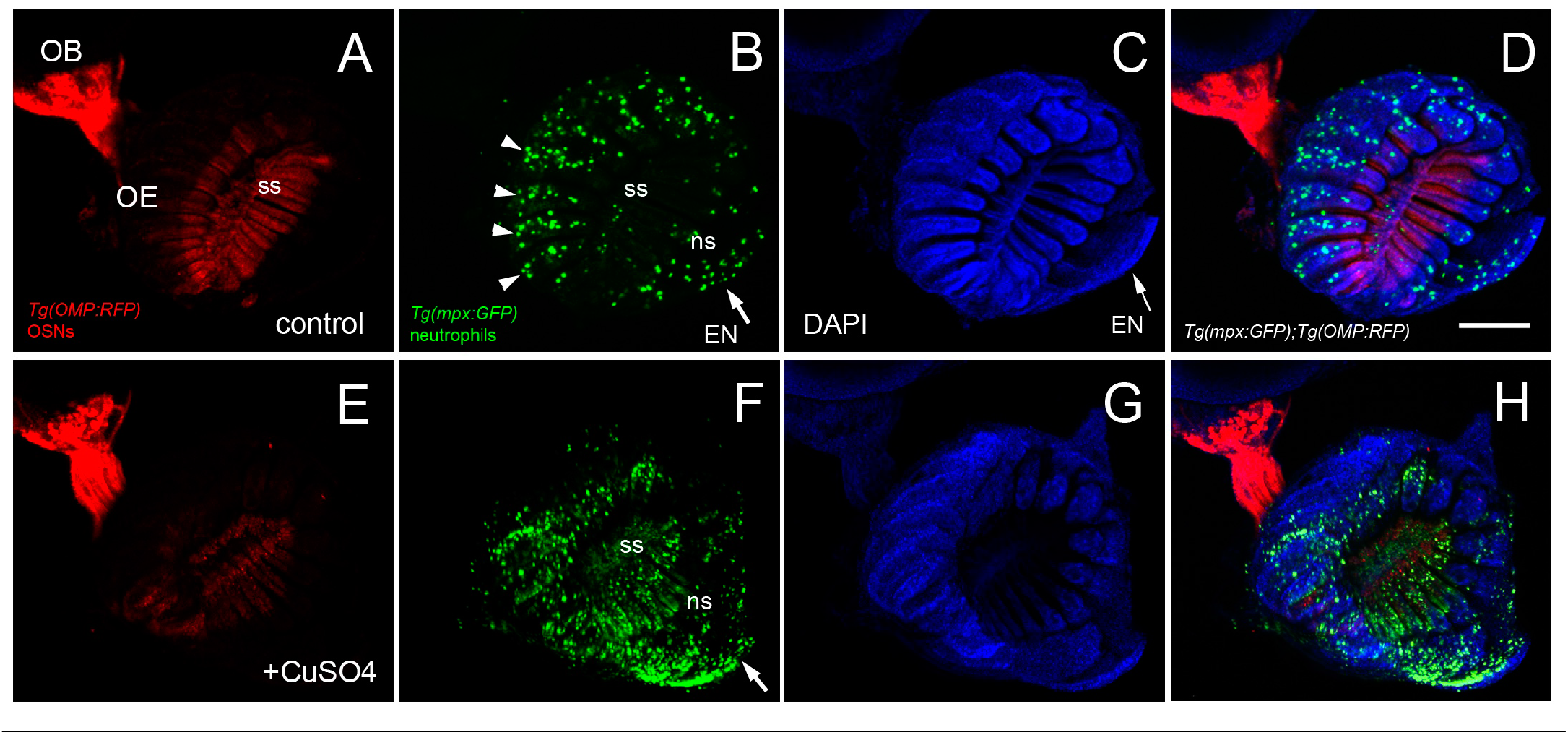
Copper exposure induces rapid increase in neutrophils in the OOs. **A**. OSNs (red) in control animal populate the sensory epithelia of the OE. **B**. Neutrophils in control animal extend up the lamellae and are found in the EN (arrow). **C.** DAPI labeling in control animal. **D.** Merge of A-C. 9 brains imaged: representative image from 1 brain. **E**. Reduced OMP:RFP labeling in OO copper exposed animals as neurons die. **F**. Increase in number of neutrophils in sensory epithelia (ss), non-sensory epithelia (ns) and EN (arrow) of in copper exposed animals. **G**. DAPI in copper exposed animals. **H**) Merge of *E-G*. 9 brains imaged: representative image from 1 brain. Scale bar = 100 µm

## Notes

### Competing Interest Statement

The authors have declared no competing interest.

